# Dinucleoside polyphosphates act as 5’-RNA caps in *Escherichia coli*

**DOI:** 10.1101/563817

**Authors:** Oldřich Hudeček, Roberto Benoni, Martin Culka, Martin Hubálek, Lubomír Rulíšek, Josef Cvačka, Hana Cahová

## Abstract

Dinucleoside polyphosphates (Np_*n*_Ns), discovered more than 50 years ago,^1^ are pleiotropic molecules present in almost all types of cells.^2^ It has been shown that their intracellular concentration can under stress conditions increase from the μM to mM range^2,3^. However, the cellular roles and mechanisms of action of Np_*n*_Ns are still speculative^4,5^. They have never been considered as part of the RNA, even though they have similar chemical structures as already known RNA caps, such as the nicotinamide adenine dinucleotide (NAD)^6–8^ and 7-methylguanylate cap^9^. Here, we show that both methylated and non-methylated Np_*n*_Ns serve as RNA caps in *Escherichia coli (E. coli).* Np_*n*_Ns are excellent substrates for T7 and *E. coli* RNA polymerases (RNAP) and efficiently initiate transcription. Further, we demonstrate that the *E. coli* decapping enzyme RNA 5’ pyrophosphohydrolase (RppH) is able to remove the Np_*n*_Ns-cap from the RNA. RppH was, however, not able to cleave the methylated forms of the Np_n_N-caps, suggesting that the methylation adds an additional layer to the RNA stability regulation. Our work introduces an original perspective on the chemical structure of RNA in prokaryotes and the function of RNA caps. This is the first evidence that small molecules like Np_*n*_Ns can act in cells via their incorporation into RNA and influence the cellular metabolism.

---

The role and chemical structure of the 5’-end of prokaryotic RNA is still unclear. The discovery of NAD^6, 10^ and Coenzyme A (CoA)^11^ as 5’ RNA caps changed the perception of the RNA structure. 5’-caps are usually cleaved by NudiX enzymes (NudC^6,12,13^, RppH^14, 15^), which can, besides their decapping role (eukaryotic Nudt16 and Dcp2^16, 17^), cleave nucleoside diphosphate linked to another moiety *(e.g.* Np_*n*_Ns^2, 18^). To investigate whether Np_*n*_Ns (Figure 1a) can serve as non-canonical initiating nucleotides (NCINs) similarly to NAD and CoA^19^, we performed *in vitro* transcription in the presence of different Np_*n*_Ns (Ap_3-6_A, Ap_4-5_G, Gp_4_G, Figure 1b) with T7 RNAP. The resulting RNA was a mixture of capped and uncapped RNA (step 1 in Figure 1b, Extended data Fig. 1a). The presence of the cap was confirmed by electrophoretic analysis after subsequent treatment with 5’-polyphosphatase and Terminator™ 5’-phosphate-dependent exonuclease (terminator, step 2 and 3 in Figure 1b). The former dephosphorylated the 5’-triphosphate RNA (5’-ppp RNA) but not any capped RNA. The terminator then digested all RNA with 5’-monophosphate termini (5’-p RNA) and left the capped RNA intact. We observed that all tested Np_*n*_Ns were excellent substrates for T7 RNAP and served as NCINs for *in vitro* transcription (Figure 1c).

**Figure 1:**
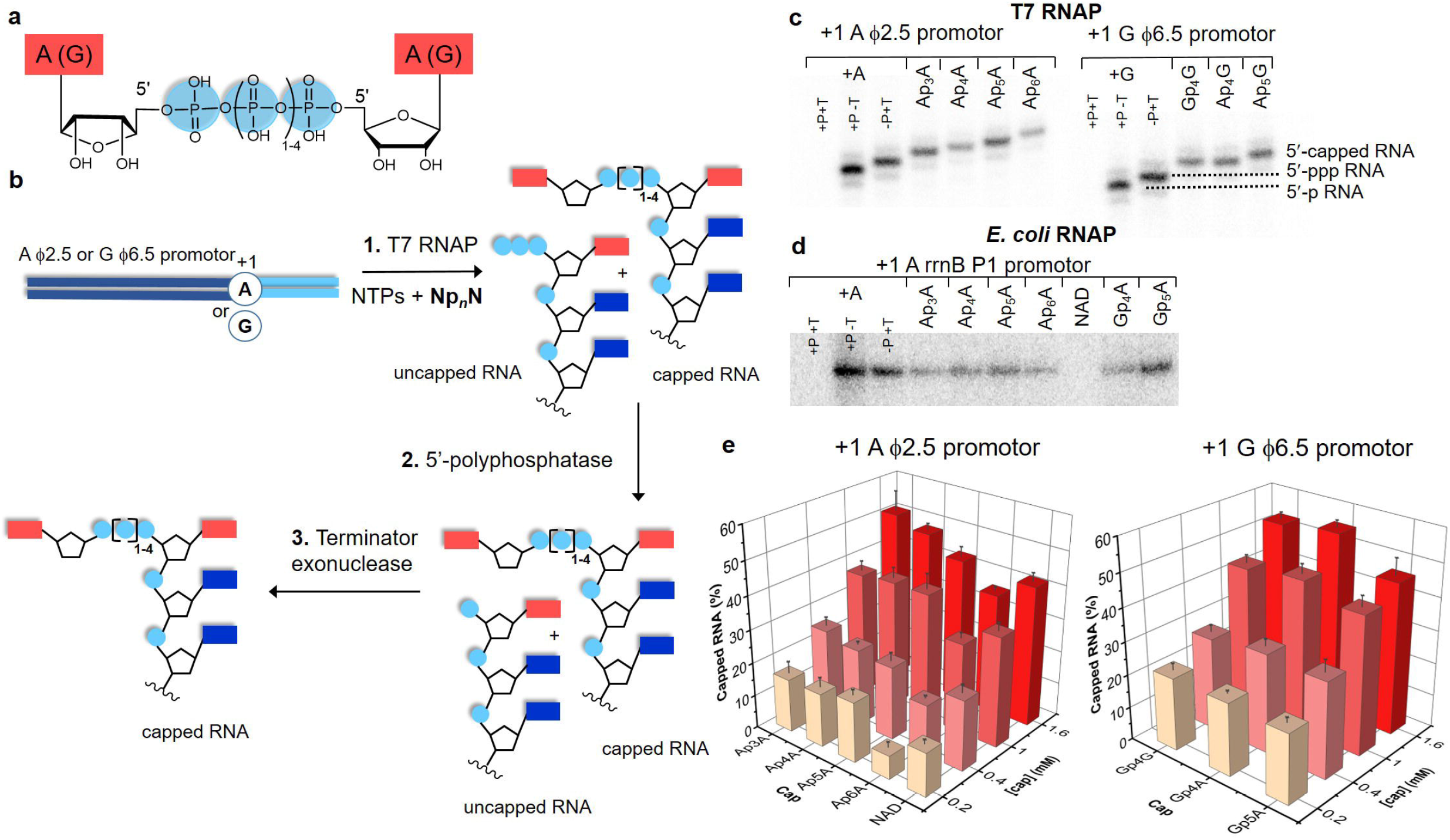
Np_*n*_Ns are excellent substrates for RNAP. **a,** The chemical structure of Np_*n*_Ns. **b,** Scheme showing the *in vitro* transcription using T7 RNAP in the presence of Np_*n*_Ns (1.6 mM) and template DNA with A φ 2.5 and G φ 6.5 promotors yielding 35mer RNA with A or G at the 5’-end. The first step resulted in a mixture of capped and uncapped RNAs, which is then treated by 5’– polyphosphatase (P) cleaving a diphosphate group from the 5’-ppp RNA leaving intact the capped RNA. In the third step the 5’-p RNA is degraded by a terminator exonuclease (T). Red (dark blue) rectangles correspond to purine base, and light blue spots depict phosphates **c,** Polyacrylamide gel electrophoretic (PAGE) analysis of the products of the *in vitro* transcription with T7 RNAP followed by P and/or T treatment. If not specified, samples were treated with both P and T enzymes. **d,** PAGE analysis of the products of *in vitro* transcription with *E. coli* RNAP and template plasmid 458 with promotor rrnB P1, which lead to the production of 144 nt long RNA having A as the first nucleotide, followed by P and/or T treatment. **e,** Percentage of different types of capped RNAs produced by *in vitro* transcription with T7 RNAP calculated from PAGE analysis. The depth axis represents various concentrations of Np_*n*_Ns (0.2, 0.4, 1 and 1.6 mM, different shades) at a constant concentration of ATP (1 mM) and GTP (1 mM). The left panel shows the percentage of Ap_3-6_A and NAD capped RNA, the right panel shows the percentage of Ap_4-5_G and Gp_4_G.

Subsequently, we tested *E. coli* RNAP that is known to accept NAD as NCIN *in vitro* and *in vivo^19–21^* (Extended data Fig. 1b). To confirm the existence of 5’-capped RNA products, we also treated them with 5’-polyphosphatase and the terminator. We found that Np_*n*_Ns are superior initiating substrates compared to NAD. The absence of NAD-capping (Figure 1d) was likely due to the different promoter sequence than previously published^19^.

To identify the best substrate for RNAP, we varied the concentrations of the Np_*n*_Ns in the presence of a constant (1 mM) ATP and GTP concentration (Figure 1e, Extended data Fig. 1c,d). The amount of capped RNA increased linearly with the concentration of Np_*n*_N. When the ratio of ATP (GTP) to Np_*n*_N was 1, we observed between 27% (for Ap_6_A) and 46% (for Ap_4_G) capped products. The NAD-capped RNA was produced in lower amounts compared to the majority of Np_*n*_Ns.

Next, we wanted to determine whether Np_*n*_Ns exist as 5’-RNA caps *in vivo* in *E. coli.* We established an LC-MS method for their detection in RNA. Because the intracellular concentration of Np_*n*_Ns is known to grow under stress conditions^2,3,22^, we collected cells in exponential (EXP, OD=0.3) and late stationary phase of growth (STA). We focused on short RNA (sRNA) where NAD-cap^10^ and CoA-cap^11^ have also been detected. The RNA was washed to remove all non-covalently interacting molecules and digested by Nuclease P1 into the form of nucleotides (Figure 2a). The negative control samples, where the addition of Nuclease P1 was omitted, did not show any signals of nucleotides or Np_*n*_Ns, which excluded the possibility of non-covalently bound contamination.

**Figure 2:**
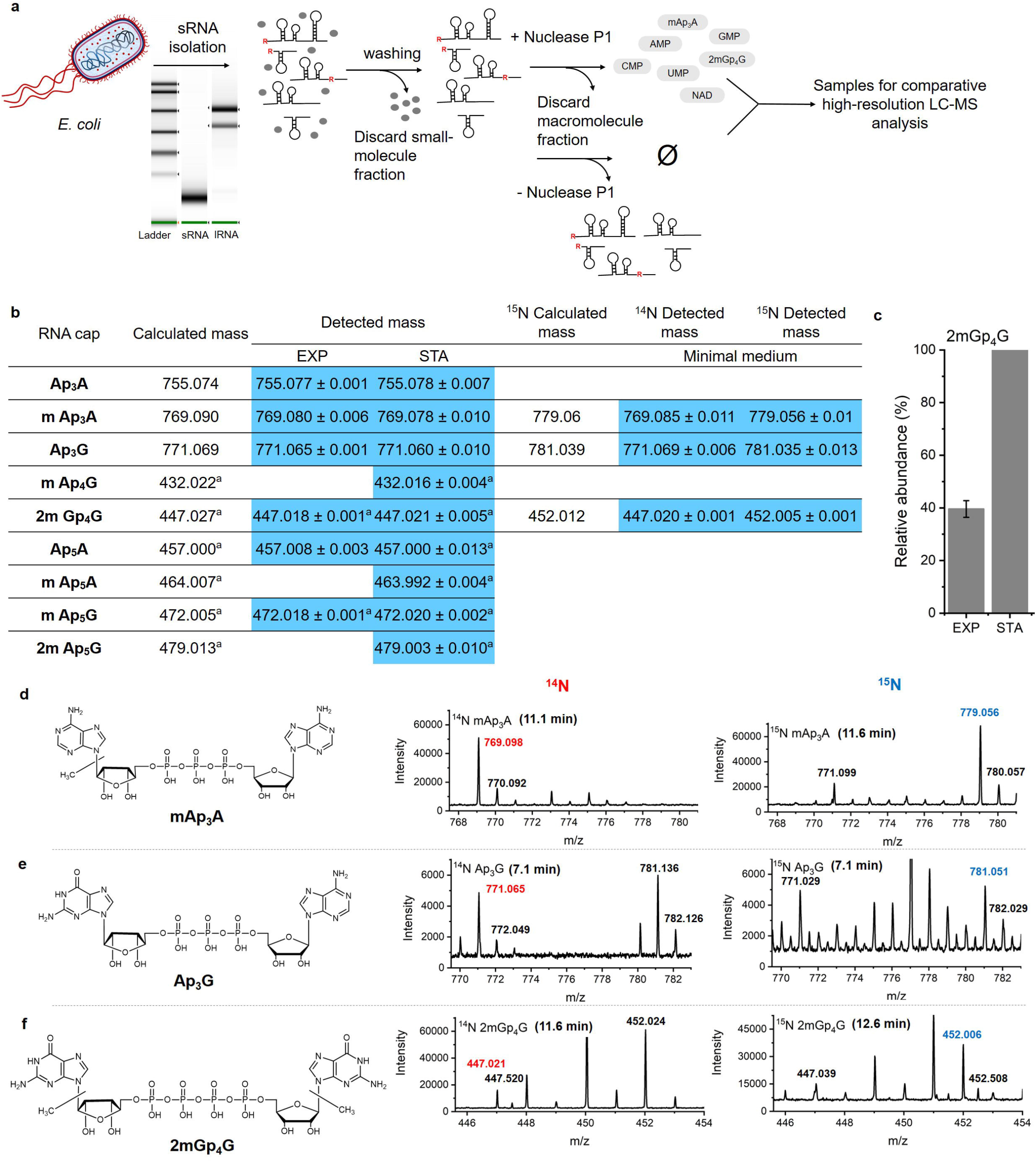
LC-MS detection of naturally occurring Np_n_N-RNA in *E. coli.* **a**, Scheme showing the RNA preparation for comparative LC-MS measurements. RNA was isolated from *E. coli* in various stages of growth. Short RNA was separated from long RNA by the RNAzol protocol and analyzed by Tape station. sRNA was separated from non-covalently bound small molecules by size exclusion chromatography and divided into two parts. One part was treated by Nuclease P1, the other was treated under identical conditions without addition of Nuclease P1 as negative control. Both samples were subjected to size exclusion chromatography again and the fraction of small molecules was analyzed by LC-MS. **b**, Table of detected *m/z* values in LC-MS analysis of digested sRNA from *E. coli* harvested in the exponential phase (EXP), the late stationary phase (STA) and after growth in minimal media with the sole source of nitrogen from ^14^NH_4_Cl (^14^N) or ^15^NH_4_Cl (^15^N). **c**, Relative quantification of dimethylated-Gp_4_G in RNA from EXP and STA growth of *E. coli.* d-f, Structures of different RNA caps and MS spectra of the detected *m/z* in RNA from *E. coli* growth in minimal media containing ^14^N or ^15^N of methyl-Ap_3_A (d, in ^14^N *m/z* 769.085 was shifted to *m/z* 779.056 in ^15^N), Ap_3_G (e, in ^14^N *m/z* 771.069 was shifted to *m/z* 781.035 in ^15^N), dimethyl-Gp_4_G (f, in ^14^N *m/z* 447.021 was shifted to *m/z* 452.006 in ^15^N).

In all the digested sRNA, we observed signals of Ap_3_A ([M-H]^-^ at *m/z* 755.077), Ap_3_G ([M-H]^-^ at *m/z* 771.065) and Ap_5_A ([M-2H]^2-^ at *m/z* 457.008) (Extended data Fig. 2a,b,c). We also observed significant signals of mono- and dimethylated forms of Np_*n*_Ns, specifically methyl-Ap_3_A ([M-H]^-^ at *m/z* 769.080), dimethyl-Gp_4_G ([M-2H]^2-^ at *m/z* 447.018) and methyl-Ap_5_G ([M-2H]^2-^ at *m/z* 472.018) (Extended data Fig. 3a,b,c). Besides the previously mentioned caps, in the STA, we detected signals of methyl-Ap4G ([M-2H]^2-^ at *m/z* 432.016), methyl-Ap_5_A ([M-2H]^2-^ at *m/z* 463.992) and dimethyl-Ap_5_G ([M-2H]^2-^ at *m/z* 479.003) (Figure 2b, Extended data Fig. 4a,b,c). We compared the amounts of dimethyl-Gp_4_G-RNA at various growth stages. The level of this cap was more than two-fold higher in the STA compared to EXP (Figure 2c). In general, a higher number of Np_*n*_Ns were detected in this phase. This may indicate that the cells in the STA lack the nutrients and methylate the Np_*n*_N-caps to preserve RNA.

To confirm the structure of the detected Np_*n*_N-caps, we grew the *E. coli* in minimal media with the sole source of nitrogen from either ^14^NH_4_Cl or ^15^NH_4_Cl. We detected only three caps: methyl-Ap_3_A, Ap_3_G, and dimethyl-Gp_4_G (Figure 2b,d,e,f Extended data Fig. 5a,b,c), because this type of growth represents another type of stress. This experiment confirmed the presence of ten nitrogen atoms in every detected molecule. To further verify the chemical structure of Np_*n*_Ns, we compared the LC-MS properties of standard Gp_4_G with the isomeric p_3_GpG. While half of the p_3_GpG was fragmented in the ionization source to p_2_GpG, the Gp_4_G stayed intact (Extended data Fig. 6a,b). The same behavior was observed for the dimethyl-Gp_4_G in the *E. coli* RNA sample, proving the internal polyphosphate chain. By linear ion trap LC-MS, we detected an intact triphosphate chain of Ap_3_A confirming its structure (Extended data Fig. 7).

Since Np_*n*_N-capped RNAs are produced in *E. coli,* degradation mechanisms of the capped RNA in *E. coli* must exist. Ap_4-6_A have been reported to be *in vitro* substrates for the *E. coli* NudiX enzyme NudH (RppH^18, 23^). RppH is an *E. coli* decapping enzyme of 5’-triphosphate^4,14,24^ and 5’-diphosphate RNA^15^. To assess whether Ap_4-6_A-capped RNA can be an RppH substrate *in vivo,* we prepared Np_*n*_N- and NAD-capped RNAs by *in vitro* transcription and tested the products as substrates for RppH. First, we added RppH to transform 5’-cappped RNA into 5’-p RNA, which then served as substrate for the terminator (Figure 3a). Electrophoretic analysis showed that Np_*n*_N-capped RNAs are cleaved into 5’-p RNA and subsequently degraded (Figure 3b,c), suggesting that Ap_4-6_A and Ap_4-5_G-capped RNAs are excellent substrates for RppH *in vitro.* However, Ap_3_A and NAD^6^ were cleaved less efficiently. Surprisingly, the Np_*n*_N-capped RNAs were always superior substrates for RppH compared to 5’-ppp RNAs. To understand the substrate specificity of RppH, we performed kinetic study of Np_*n*_N-capped RNAs. The decapping reaction of Ap_*4,5*_N RNA was almost quantitative within 5 min while the corresponding 5’-ppp RNA was decapped only by 50% within 40 min (Figure 3d,e, Extended data Fig 8).

**Figure 3:**
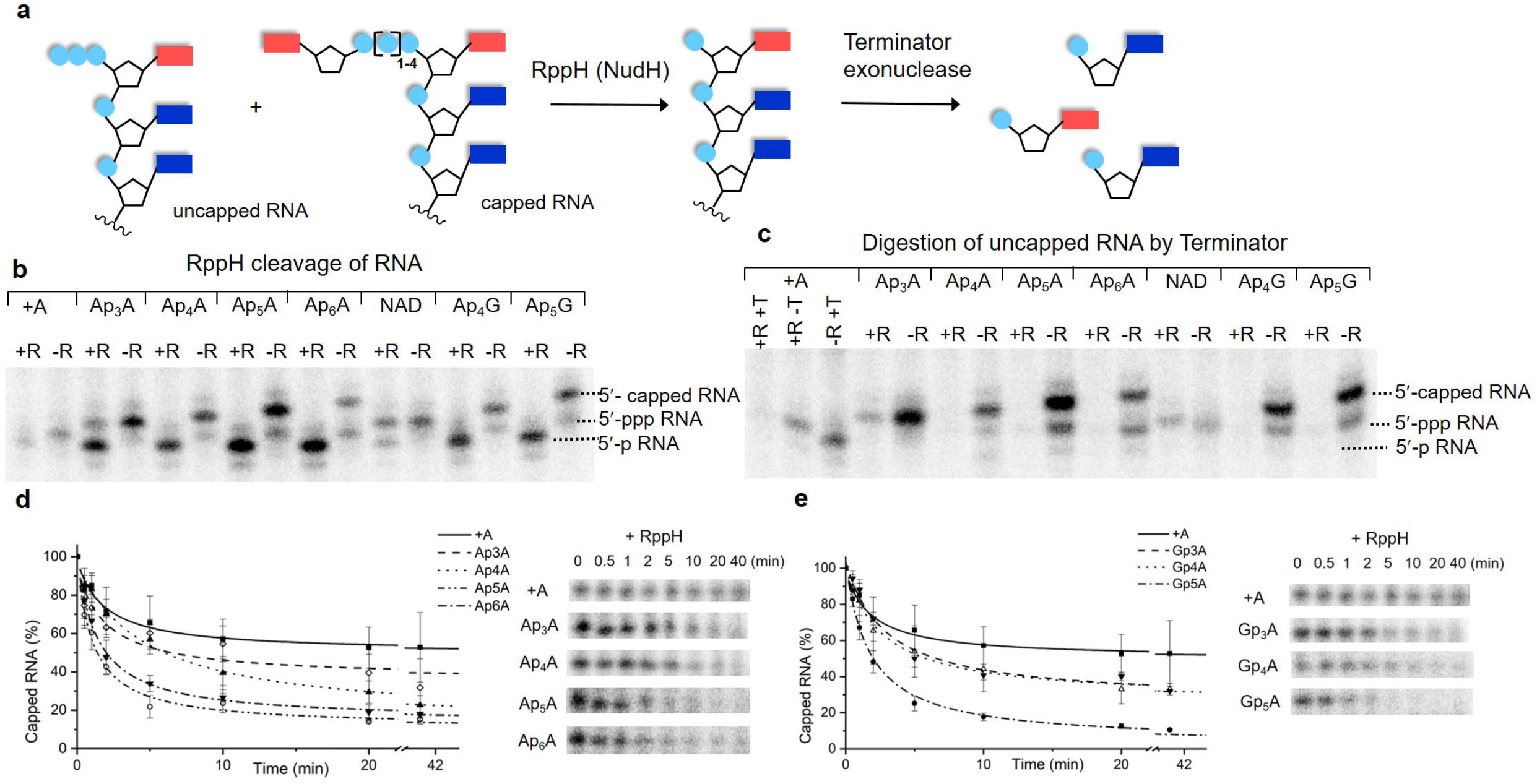
RppH cleavage of Np_n_N-capped RNA. **a,** Scheme showing the degradation of capped and uncapped RNA from *in vitro* transcription using RppH and terminator exonuclease. **b,** PAGE analysis of *in vitro* transcribed RNA treated with RppH (+R) or without RppH (-R). **c,** PAGE analysis of RNA from **b** after the reaction with the terminator (T). **d,e,** Kinetic studies of RppH cleavage of Np_n_N-capped and uncapped RNA stopped after 30 s, 1, 2, 5, 10, 20 and 40 min and analysed by PAGE. **d,** Ap_n_A-RNAs in comparison with pppA-RNA. **e,** Gp_n_A-RNAs in comparison with pppA-RNA.

As Np_*n*_N RNAs are excellent substrates for RppH *in vitro,* we constructed a mutant *E. coli* strain KS47 *(Δrpph)* with knock-down RppH to enhance the *in vivo* concentration of Np_*n*_N RNAs. The RNA isolated from the mutant strain did not show any significant difference in Np_*n*_N-capping compared to wild type, possibly due to the redundancy of NudiX enzymes.^25^

To reveal the effect of Np_*n*_N-caps methylation on the RNA, we performed molecular dynamics (MD) simulations of the interaction between RppH and Np_*n*_Ns-capped RNA (Gp_4_G-G and 2mGp_4_G-G). The LC-MS analysis confirmed the presence of one methyl group per guanosine moiety in 2mGp_4_G (Extended data Fig. 9). We speculated that these methylations could be in the positions N^7^ (m^7^G) and O2’ (Gm) similarly to eukaryotic RNA caps. We observed in MD simulations that the interactions with the arginines R28 and R86 were lost when the methylations were present (Figure 4a). These arginines are responsible for purines binding via cation-π stacking. This finding demonstrates that methylation of Np_*n*_Ns-caps in RNA can hamper decapping by RppH.

**Figure 4:**
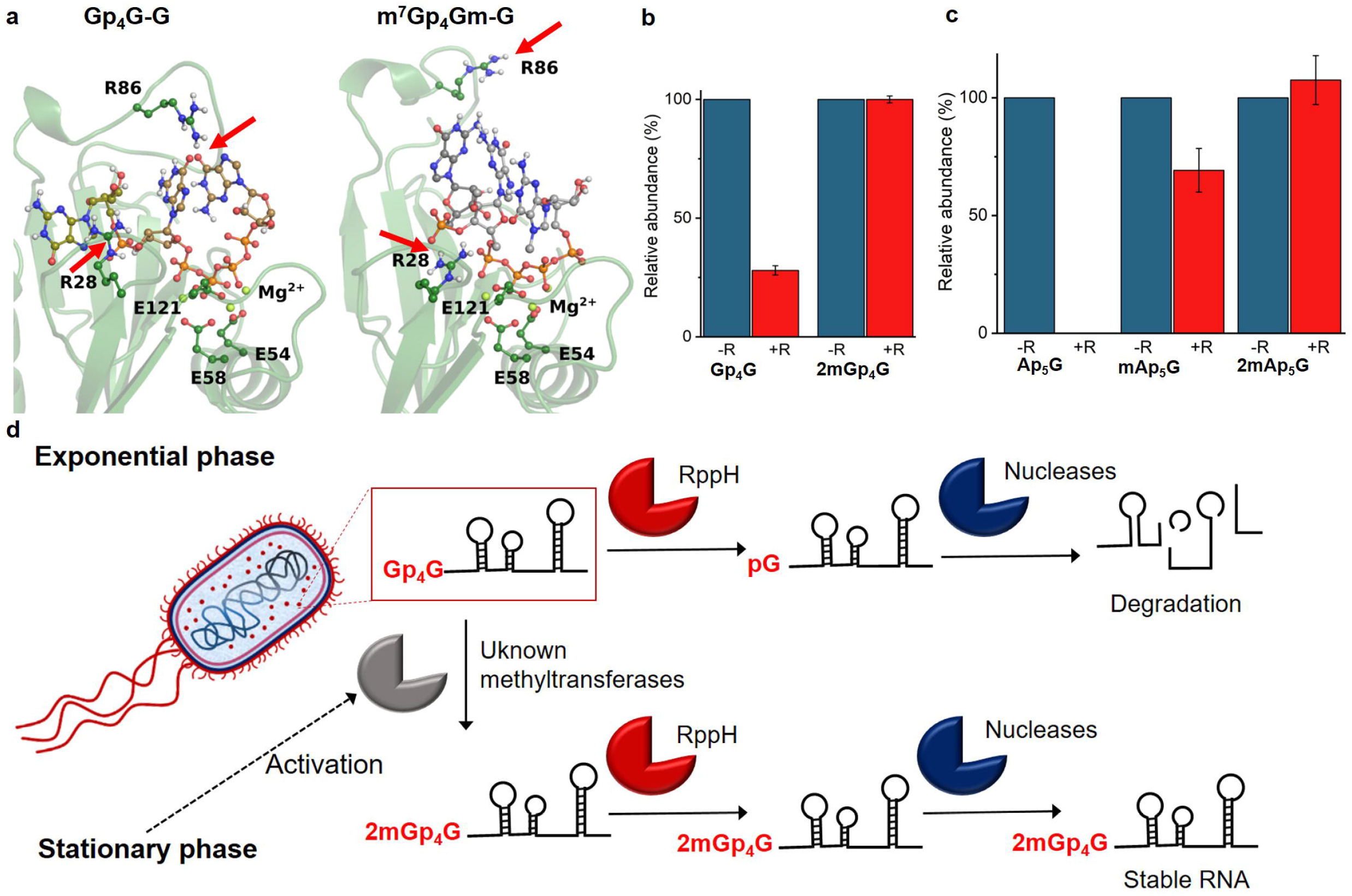
Role of RppH in the cleavage of Np_n_N-capped RNA in *E. coli.* **a,** Snapshots from molecular dynamics simulation of the interaction of RppH with Gp_4_G-G and m^7^Gp_4_Gm-G after 200 ns. **b,c** Relative abundance of non-methylated Np_*n*_Ns (left) and methylated Np_*n*_Ns (right) as derived from EIC in the sRNA fraction spiked with Gp_4_G-RNA before (blue) and after 1 h RppH treatment (red) **b** and spiked with Gp_5_A-RNA before (blue) and after 1 h RppH treatment (red) **c. d,** Hypothetic cellular processing of RNA in *E. coli* at different stages of growth.

To experimentally prove the MD findings, we added RppH into the mixture of isolated sRNA with a spiked model Gp_4_G-RNA or Gp_5_A-RNA to compare the activity of RppH on methylated and non-methylated substrates. We found that majority of the model capped RNAs were cleaved within 1 h while the amount of naturally present methyl-Ap_5_G-RNA slightly decreased and dimethyl-Gp_4_G/Ap_5_G-RNA remained unchanged (Figure 4b,c) This confirms that the methylation stabilizes the Np*n*N capped RNAs against cleavage by RppH.

In summary, we identified Np_*n*_Ns as novel 5’-RNA caps, which are incorporated into RNA by RNAPs. We found that Np_*n*_N RNAs were cleaved by the *E. coli* RppH decapping enzyme. Caps with long polyphosphate chains were cleaved most efficiently and are better substrates than 5’-ppp RNAs. This suggests that they may be the main cellular targets of RppH. In the cell, we detected the presence of Np_*n*_Ns in sRNA, including their methylated variants. MD simulations and experiments revealed that the methylation protects the caps from the RppH cleavage. The amounts of caps and their methylations increased in the STA. Hence, we propose that bacteria use methylated caps to stabilize some RNAs under stress (Figure 4d). In the exponential phase, the metabolism is efficient and the turnover of the macromolecules is fast. Therefore, the methylated caps are not necessary and were detected in low amount. In the stationary phase, cells lack nutrients so they need to find a strategy to save macromolecules. The methylation of the Np_*n*_N-caps can be the way to preserve important RNA molecules. It is intriguing to conceive that many functions of Np_*n*_Ns can be explained via their RNA capping potential. Moreover, we show that RNA possesses at its 5’-termini previously unknown structures that may interact with a wide range of cellular partners and influence *e.g.* cellular reaction to starvation. Besides searching for methyltransferases responsible for the methylation of the Np_*n*_N-caps, the biggest challenges lie in the development of specific techniques for identification of Np_*n*_N-capped RNAs.

**Supplementary Information** is available in the online version of the paper

## Acknowledgements

We thank Dr. K. Stříšovský for help with the preparation of mutant *E. coli* strain KS47 (*Δrpph*), Dr. L. Krásný for critical reading of the manuscript and advices, Dr. P. Reyes-Gutierrez and other members of Cahova lab for their help and discussion. This work was supported by the Ministry of Education, Youth and Sports (Czech Republic), program ERC CZ (LL1603).

## Author Contributions

O.H., R.B., M.H. and H.C. conceived the study, designed the experiments. O.H. and R.B. performed the experiments. J.C. L.R. and H.C. supervised the work. M.C. performed molecular dynamics study. O.H., R.B., M.C. and H.C. wrote the paper.

## Author Information

The authors declare no competing financial interests. Correspondence and requests for materials should be addresses to H.C.

## References

1 Zamecnik, P. G., Stephenson, M. L., Janeway, C. M. & Randerath, K. Enzymatic synthesis of diadenosine tetraphosphate and diadenosine triphosphate with a purified lysyl-sRNA synthetase. Biochemical and Biophysical Research Communications 24, 91–97, doi:https://doi.org/10.1016/0006-291X(66)90415-3 (1966).

2 McLennan, A. G. Ap4a and Other Dinucleoside Polyphosphates. (Taylor & Francis, 1992).

3 VanBogelen, R. A., Kelley, P. M. & Neidhardt, F. C. Differential induction of heat shock, SOS, and oxidation stress regulons and accumulation of nucleotides in Escherichia coli. Journal of Bacteriology 169, 26–32, doi:10.1128/jb.169.1.26-32.1987 (1987).

4 McLennan, A. G. Dinucleoside polyphosphates—friend or foe? Pharmacology & Therapeutics 87, 73–89, doi:https://doi.org/10.1016/S0163-7258(00)00041-3 (2000).

5 Despotović, D. et al. Diadenosine tetraphosphate (Ap4A) – an E. coli alarmone or a damage metabolite? The FEBS Journal 284, 2194–2215, doi:10.1111/febs.14113 (2017).

6 Cahová, H., Winz, M.-L., Höfer, K., Nübel, G. & Jäschke, A. NAD captureSeq indicates NAD as a bacterial cap for a subset of regulatory RNAs. Nature 519, 374, doi:10.1038/nature14020 https://www.nature.com/articles/nature14020#supplementary-information (2014).

7 Walters, R. W. et al. Identification of NAD+ capped mRNAs in Saccharomyces cerevisiae. Proc. Natl. Acad. Sci. U. S. A. 114, 480–485, doi:10.1073/pnas.1619369114 (2017).

8 Jiao, X. et al. 5’ End Nicotinamide Adenine Dinucleotide Cap in Human Cells Promotes RNA Decay through DXO-Mediated deNADding. Cell 168, 1015–1027.e1010, doi:https://doi.org/10.1016/j.cell.2017.02.019 (2017).

9 Adams, J. M. & Cory, S. Modified nucleosides and bizarre 5’-termini in mouse myeloma mRNA. Nature 255, 28–33, doi:10.1038/255028a0 (1975).

10 Chen, Y. G., Kowtoniuk, W. E., Agarwal, I., Shen, Y. & Liu, D. R. LC/MS analysis of cellular RNA reveals NAD-linked RNA. Nature Chemical Biology 5, 879, doi:10.1038/nchembio.235 https://www.nature.com/articles/nchembio.235#supplementary-information (2009).

11 Kowtoniuk, W. E., Shen, Y., Heemstra, J. M., Agarwal, I. & Liu, D. R. A chemical screen for biological small molecule–RNA conjugates reveals CoA-linked RNA. Proceedings of the National Academy of Sciences 106, 7768–7773, doi:10.1073/pnas.0900528106 (2009).

12 Höfer, K. et al. Structure and function of the bacterial decapping enzyme NudC. Nature Chemical Biology 12, 730, doi:10.1038/nchembio.2132 https://www.nature.com/articles/nchembio.2132#supplementary-information (2016).

13 Zhang, D. et al. Structural basis of prokaryotic NAD-RNA decapping by NudC. Cell Research 26, 1062, doi:10.1038/cr.2016.98 https://www.nature.com/articles/cr201698#supplementary-information (2016).

14 Deana, A., Celesnik, H. & Belasco, J. G. The bacterial enzyme RppH triggers messenger RNA degradation by 5’ pyrophosphate removal. Nature 451, 355, doi:10.1038/nature06475 https://www.nature.com/articles/nature06475#supplementary-information (2008).

15 Luciano, D. J., Vasilyev, N., Richards, J., Serganov, A. & Belasco, J. G. A Novel RNA Phosphorylation State Enables 5’ End-Dependent Degradation in Escherichia coli. Molecular Cell 67, 44–54.e46, doi:https://doi.org/10.1016/j.molcel.2017.05.035 (2017).

16 Grudzien-Nogalska, E. & Kiledjian, M. New insights into decapping enzymes and selective mRNA decay. Wiley Interdisciplinary Reviews: RNA 8, e1379, doi:doi:10.1002/wrna.1379 (2017).

17 Song, M.-G., Li, Y. & Kiledjian, M. Multiple mRNA Decapping Enzymes in Mammalian Cells. Molecular Cell 40, 423–432, doi:10.1016/j.molcel.2010.10.010.

18 McLennan, A. G. The Nudix hydrolase superfamily. Cellular and Molecular Life Sciences CMLS 63, 123–143, doi:10.1007/s00018-005-5386-7 (2006).

19 Bird, J. G. et al. The mechanism of RNA 5’ capping with NAD+, NADH and desphospho-CoA. Nature 535, 444, doi:10.1038/nature18622 https://www.nature.com/articles/nature18622#supplementary-information (2016).

20 Barvík, I., Rejman, D., Šanderová, H., Panova, N. & Krásný, L. Non-canonical transcription initiation: the expanding universe of transcription initiating substrates. FEMS Microbiology Reviews 41, 131–138, doi:10.1093/femsre/fuw041 (2016).

21 Vvedenskaya, I. O. et al. CapZyme-Seq Comprehensively Defines Promoter-Sequence Determinants for RNA 5’ Capping with NAD+. Molecular Cell 70, 553–564.e559, doi:https://doi.org/10.1016/j.molcel.2018.03.014 (2018).

22 Lee, P. C., Bochner, B. R. & Ames, B. N. AppppA, heat-shock stress, and cell oxidation. Proceedings of the National Academy of Sciences 80, 7496–7500, doi:10.1073/pnas.80.24.7496 (1983).

23 Bessman, M. J. et al. The gene, ygdP, associated with the invasiveness of Escherichia coli K1, designates a nudix hydrolase (Orf 176) active on Adenosine (5’) pentaphospho (5’) adenosine. Journal of Biological Chemistry, doi:10.1074/jbc.M107032200 (2001).

24 Foley, P. L., Hsieh, P.-k., Luciano, D. J. & Belasco, J. G. Specificity and Evolutionary Conservation of the Escherichia coli RNA Pyrophosphohydrolase RppH. Journal of Biological Chemistry, doi:10.1074/jbc.M114.634659 (2015).

25 McLennan, A. G. Substrate ambiguity among the nudix hydrolases: biologically significant, evolutionary remnant, or both? Cellular and Molecular Life Sciences 70, 373–385, doi:10.1007/s00018-012-1210-3 (2013).

