## Supplementary material for "Dinucleoside polyphosphates act as 5’-RNA caps in *Escherichia coli*": Methods

### General

All reagents were either purchased from Merck or Jena Biosciences and used without further purification. Oligonucleotides were purchased from Generi Biotech.

Denaturating polyacrylamide gels (PAGE) were visualized by a Typhoon FLA 9500 imaging system.

### *In vitro* transcription with T7 RNAP

*In vitro* transcription was performed using a modification of a previously published method[1] in a 25 $\mu$L mixture containing: 80 ng/$\mu$L of template DNA (35A: 5'-CAGTAATACGACTCACTATT**A**GGGAAGCGGGCATGCGGCCAGCCATAGCCGATCA or 35G: 5'-CAGTAATACGACTCACTATA**G**GGGAAGCGGGCATGCGGCCAGCCATAGCCGATCA), 1 mM UTP, 1 mM CTP, 0.8 mM GTP or 0.8 mM ATP, respectively, and 0.2 $\mu$L $\alpha$ $^{32}$P GTP or $\alpha$ $^{32}$P ATP (activity: 9.25 MBq in 25 $\mu$L), respectively, 0.2-1.6 mM Np$_n$Ns, 5% DMSO, 0.12% triton X-100, 12 mM DTT, 4.8 mM magnesium chloride and 10x reaction buffer for T7 RNAP (40 mM Tris-HCl, 6 mM $MgCl_2$, 1 mM DTT, 2 mM spermidine, pH 7.9 at 25°C) and 62.5 units of T7 RNAP. The mixture was incubated for 2 h at 37°C. The samples (3 $\mu$L) were mixed with 2x RNA loading dye (New England BioLabs, NEB) and analyzed by 12% PAGE (600 V, 3.5 h).

### DNAse I treatment

The DNA template was digested by DNAse I to obtain pure RNA. 25 $\mu$L of the transcription mixture, 3 $\mu$L of 10x reaction buffer for DNAse I (10 mM Tris-HCl, 2.5 mM $MgCl_2$, 0.5 mM $CaCl_2$, pH 7.6 at 25°C, supplied with the enzyme) and 4 units of DNAse I (NEB) were incubated at 37°C for 60 min. The enzyme was heat deactivated at 75°C for 10 min followed by immediate cooling on ice. All samples were purified on RNA mini Quick Spin Columns (Merck) for further use.

### RNA 5′-polyphosphatase treatment

2.5-3 $\mu$g of RNA (10 $\mu$L) were treated with 20 units of 5′-polyphosphatase (Epicentre) in the solution of 10x buffer (0.05 M HEPES-KOH, pH 7.5, 0.1 M NaCl, 1.0 mM EDTA, 0.1% β-mercaptoethanol, and 0.01% Triton X-100) in a total volume of 20 $\mu$L for 1 h at 37°C. For the negative control, the enzyme was replaced by the same amount of water. The samples (3 $\mu$L) were mixed with 2x RNA loading dye and analyzed by 12% PAGE (600 V, 3.5 h).

### Terminator™ 5´-Phosphate-Dependent Exonuclease treatment

Because of the incompatibility of the buffers, all samples were purified on RNA mini Quick Spin Columns before the reaction. The RNA (500 ng) was treated with 1 unit of Terminator™ 5´-Phosphate-Dependent Exonuclease (Epicentre) in the solution of 10x reaction buffer A before the mixture was incubated at 30°C for 1 h.

### *E. coli* RNA polymerase in vitro transcription

*In vitro* transcription was performed in a volume of 40 $\mu$L. The first part of the mixture contained 10 ng/$\mu$L of template plasmid 458 with promotor rrnB P1 (5'GAATTCGCGGTCAGAAAATTATTTTAAATTTCCTCTTGTCAGGCCGGAATAACTCCCTATAATGCGCCACCACTG ACACG**A**AGCTTGGGTCCCACCTGACCCCATGCCGAACTCAGAAGTGAAACGCCGTAGCGCCGATGGTAGTGTGGG

GTCTCCCCATGCGAGAGTAGGGAACTGCCAGGCATCAAATAAAACGAAAGGCTCAGTCGAAAGACTGGGCCTTT), 8 μL of 5x buffer (supplied with the enzyme), 90 mM NaCl, 0.2 mM ATP, 0.8 mM $Np_nNs$ and 1 unit of *E. coli* RNA polymerase holoenzyme (NEB). This mixture was pre-incubated for 10 min at 37°C. The transcription was started by addition of an initiation mixture containing 0.2 mM CTP and UTP, 0.15 mM GTP and 1 μL $\alpha$ $^{32}$P GTP (activity: 9.25 MBq in 25 μL) and incubated at 37°C for 1 h.

**Calculations of capped RNA**

The experiments were performed in triplicates and the amount of capped RNA was calculated from PAGE analysis using the software ImageJ[2]. The sum of areas under the peaks corresponding to the 5′-triphosphate RNA ($Ar_p$) and 5′-capped RNA ($Ar_{cap}$) were integrated and regarded as 100%. The percentage of 5′-capped RNA species were calculated according to: $(Ar_{cap}/ Ar_p + Ar_{cap})\cdot 100$.

**Cleavage of capped RNA by RppH**

To test the cleavage of the 5′-caps, the RNA samples were divided into two parts. The positive control contained 700 ng of the RNA (*in vitro* transcription, DNAse I treatment, purified on RNA mini Quick Spin Columns), 2 μL of 10x buffer 2 (500 mM NaCl, 100 mM Tris-HCl, 100 mM $MgCl_2$, 10 mM DTT, pH 7.9 at 25°C, supplied with the enzyme) and 5 units of RppH (NEB). Water was used as the negative control. The mixtures were incubated at 37°C for 1 h and purified on RNA mini Quick Spin Columns. 3 μL of the purified samples were mixed with 2x RNA loading dye for analysis and 6 μL were used for the Terminator treatment.

**Kinetic studies**

For the kinetic studies, RNA samples after *in vitro* transcription with $^{32}$P GTP, DNase I treatment and purification on RNA mini Quick Spin Columns were used. The 20 μL mixture contained 2400 ng of the studied RNA, buffer 2 and 0.2 units of RppH enzyme. The mixture was incubated at 37°C, aliquots of 2 μL were collected at 0, 0.5, 1, 2, 5, 10, 20, and 40 min and mixed immediately with 2 μL of 2x RNA loading dye. All samples were analyzed by 12% PAGE. The experiments were performed in triplicates and the amount of capped RNA was calculated from PAGE analysis using the software ImageJ. The areas under the peaks corresponding to 5′-triphosphate RNA or 5′-capped RNA were integrated and plotted against time by regarding the area at 0 min as 100%.

***E. coli* growth condition**

The *E. coli* strain DH5α containing pUC18 was used for studying RNA modification in three different growth conditions: exponential phase, stationary phase and heat shock. The growths were performed in LB broth (Merck) in the presence of 0.1 mg/mL of Ampicillin (Amp, Merck). Parallel cultures of *E. coli* were inoculated in Erlenmeyer flasks from cultures grown on LB Agar. Cultures (1 L) were grown at 37°C until an optical density at 600 nm ($OD_{600}$) of 0.3 was reached (Exponential phase). Cultures (1 L) were harvested after 2 hours they reached an $OD_{600}$ of 1.6 (Stationary phase). The heat shock experiment consisted of two phases. In the first phase, cells (1 L) were grown at 28°C until an $OD_{600}$ of 0.3 was reached. This was followed by a second phase, in which the temperature was held at 50°C for 1.5 hour. All cells were harvested by centrifugation at 5,000 g for 10 min. The pellets were washed once with PBS and stored at -80°C.

**$^{14}$N/$^{15}$N isotope RNA labelling**

*E. coli* strains DH5α containing pUC18 were grown in M9 minimal medium containing 1x M9 salts (5x concentrate: 34 g/L $Na_2HPO_4$, 29 g/L of $KH_2PO_4$, 2.5 g/L of NaCl, 5 g/l $NH_4Cl$), 2mM $MgSO_4$, 0.1 mM $CaCl_2$ and 0.4% glucose). The growth was performed in the presence of 0.1 mg/mL of Amp. Parallel cultures (1 L each) containing $^{14}NH_4Cl$ or $^{15}NH_4Cl$ (as sole source of nitrogen) were inoculated in Erlenmeyer flasks from cultures grown overnight in M9 medium (50 mL). All cells were harvested by centrifugation at 5,000 g for 10 min after 48 h of growth (OD=0.5), washed once with PBS and stored at -80°C.

**Mutant *E. coli* strain KS47 (Δ*rpph)***

The *E. coli* strain KS47 was used for construction of a mutant with knock-down *rpph* gene (Δ*rpph*) using the protocol for one-step inactivation of chromosomal genes [3]. The Δ*rpph* and wild type (wt) were transfected with pUC18 using the electroporation technique and grown in LB medium. The cultures were harvested in the exponential and late stationary phases of growth by centrifugation at 5,000 g for 10 min. The pellets were washed once with PBS and stored at -80°C.

**Isolation and purification of short RNA**

Short RNA from *E. coli* was isolated using the RNAzol protocol (Merck). The pellets (stemming from 1 L culture of exponential and heat shock growth, respectively, and from 0.5 L stationary phase growth) were suspended in 4 mL lysozyme solution (1 mg/mL, Merck) each and incubated for 1 h at 7°C. RNAzol (10 mL) was added to the lysate to isolate the RNA. After shaking vigorously for 15 s, the mixture was incubated for 15 min on ice and centrifuged at 6,000 g for 45 min. In this step, the DNA, proteins and most polysaccharides formed a semisolid pellet at the bottom of the tube. The RNA, which remained in the supernatant, was mixed with 0.4 volumes of 75% ethanol (v/v). After storing on ice for 10 min, the sample was centrifuged at 6,000 g for 16 min for pelleting the long RNA. The supernatant containing short RNA was mixed with 0.8 volumes of isopropanol. After storing on ice for 30 min, the sample was centrifuged at 6,000 g for 40 min for pelleting the short RNA. The pellets containing the long and short RNA were washed twice with 75% ethanol (v/v) and 70% isopropanol (v/v), respectively. Both pellets were dissolved in small amounts of water and stored at -80°C. To remove possible contaminants, all short RNA samples were submitted to size exclusion chromatography for five times using an Amicon Ultra 3 KDa cut-off (Merck). All experiments were performed in at least triplicates.

**RppH experiments with sRNA**

sRNA (1 mg) from the stationary phase was spiked with 10 μg of model $Gp_4G$-RNA or $Ap_5G$-RNA and divided into two halves. The first part was diluted to a final volume of 300 μL with 30 μL of 10x buffer 2 (500 mM NaCl, 100 mM Tris-HCl, 100 mM $MgCl_2$, 10 mM DTT, pH 7.9 at 25°C, supplied with the enzyme) and 15 U of RppH (NEB). The second half was used as a negative control and also diluted to a final volume of 300 μL using buffer without the addition of RppH. The mixtures were incubated at 37°C for 1 h, and further used for LC-MS analysis.

**sRNA digestion for LC-MS**

1 mg of short RNA was divided into two aliquots; one aliquot of 0.5 mg was digested by 10 U of Nuclease P1 (Merck) in 50 mM ammonium acetate buffer (pH 4.5) at 37°C for 1 h. The second aliquot of 0.5 mg was incubated without enzyme and used as negative control. The digest was purified over Amicon-Millipore filters 10 kDa (Merck). The flow-through was dried up on a Speedvac system and dissolved in 10 μL of a

mixture of acetonitrile (10%) and ammonium acetate (10 mM, pH 12). The final pH was adjusted to 10 using a solution of NaOH.

**Preparation of model Np$_n$N-capped RNA for LC-MS**

*In vitro* transcription was performed in the 25 μL mixture containing 80 ng/μL of template DNA (35A or 35G), 1 mM UTP, 1 mM CTP, 1 mM GTP, 1 mM ATP, 1.6 mM Np$_n$Ns, 5% DMSO, 0.12% triton X-100, 12 mM DTT, 4.8 mM magnesium chloride and 10x reaction buffer for T7 RNAP (40 mM Tris-HCl, 6 mM $MgCl_2$, 1 mM DTT, 2 mM spermidine, pH 7.9 at 25°C) and 62.5 units of T7 RNAP. The mixture was incubated for 2 h at 37°C. Afterwards, the solutions were treated by DNAse I and purified on an RNA mini Quick Spin Column. The samples were digested by Nuclease P1 and analyzed by LC-MS.

***In vitro* transcription with T7 RNAP for synthesis of standard pppGpG**

*In vitro* transcription was performed in the 50 μL mixture containing 120 ng/μL of template DNA (2ntG: 5'-CAGTAATACGACTCACTATA**G**G), 2.0 mM GTP, 5% DMSO, 0.12% triton X-100, 12 mM DTT, 4.8 mM magnesium chloride and 10x reaction buffer for T7 RNAP (40 mM Tris-HCl, 6 mM $MgCl_2$, 1 mM DTT, 2 mM spermidine, pH 7.9 at 25°C) and 125 units of T7 RNAP. The mixture was incubated for 2 h at 37°C. The mixture was filtered on an Amicon Ultra 0.5 Centrifugal Filter 3kDa (Merck), which was then washed with water (2x 150 μL). The combined flow through was evaporated on a Speedvac system.

**LC-MS Data Collection and Analysis**

LC-MS was performed using a Waters Acquity UPLC SYNAPT G2 instrument with an Acquity UPLC BEH Amide column (1.7 µm, 2.1 mm x 150 mm, Waters). The mobile phase A consisted of 10 mM ammonium acetate, pH 9, and the mobile phase B of 100% acetonitrile. The flow rate was kept at 0.25 mL/min and the mobile phase composition gradient was as follows: 80% B for 2 min; linear decrease to 68.7% B over 13 min; linear decrease to 5% B over 3 min; maintaining 5% B for 2 min; returning linearly to 80% B over 2 min. For the analysis, electrospray ionization (ESI) was used with a capillary voltage of 1.80 kV, a sampling cone voltage of 20.0 kV, and an extraction cone voltage of 4.0 kV. The source temperature was 120°C and the desolvation temperature 550°C, the cone gas flow rate was 50 L/h and the desolvation gas flow rate 250 L/h. The detector was operated in negative ion mode. For each sample, 8 µL of the dissolved material was injected.

Triplicates of Nuclease P1-digested RNA samples were used to identify Np$_n$Ns. Ions with less than 50 counts were not considered for further analysis. MassLynx software was used for data analysis and quantification of the relative abundance of dimethyl-Gp$_4$G.

**MS$^3$ Fragmentation Analysis**

The fragmentations studies were performed using SCIEX QTRAP 6500+ instrument with an Acquity UPLC BEH Amide column (1.7 µm, 2.1 mm x 150 mm, Waters). Mobile phase A was 10 mM ammonium acetate pH 9, and mobile phase B was 100 % acetonitrile. The flow rate was a constant 0.25 mL/min and the mobile phase composition was as follows: 80 % B for 2 min; linear decrease over 3 min to 50 % B; maintain at 50 % B for 1 min before returning linearly to 80 % B over 2 min. Electrospray ionization (ESI) was used with curtain gas of 35 (arbitrary units), Ionspray voltage of 4.5 kV. The ion source gas was 50 (arbitrary units), and the drying gas temperature was 400°C. The declustering potential was -200 V, the entrance potential -10 V, collision energy - 46 V, excitation energy 0.1 V. The detector was operated in negative ion mode. For the confirmation of $Ap_3A$ structure, the first precursor ion was $[M - H]^-$ at *m/z* 754.96 and as second ion the ATP fragment was selected $[M - H]^-$ at *m/z* 487.8. For each sample, 8 µL of the dissolved material was injected.

**Molecular dynamics (MD) simulations**

All models were based on a crystal structure of *E. coli* RppH in complex with modified ppcpAG 5′-capped dinucleotide (PDB ID 4S2Y). We modified the adenine to guanine and extended the 5′ nucleotide triphosphate into $Gp_4G$ or the methylated $m^7Gp_4Gm$. Hydrogens were added so that the amino acids were present in the standard protonation state at pH 7. Three catalytic magnesium ions were kept in place. The complex was solvated in an 83 x 68 x 71 Å box of water molecules. Some water molecules were replaced by sodium and chlorine ions to neutralize the complex charge and mimic the cellular ionic strength (0.15M). The MD simulations were performed using the NAMD software[4] and AMBER force field with ff14SB[5] parameters for the protein, OL3[6] parameters for RNA, the SPC/E model of water[7] and corresponding parameters for monovalent ions [8] and magnesium[9]. Parameters for non-canonical RNA caps were constructed using existing RNA parameters and available parameters for polyphosphates[10]. Partial atomic charges were fitted based on quantum-chemical HF/6-31G* calculations using Gaussian 09 [11]using the RESP procedure AmberTools[12] [10]. We used periodic boundary conditions in the MD simulations to emulate a bulk solvent. An NPT scheme with Langevain temperature and pressure control was applied. The following equilibration protocol was used: (1) 1000 steps of conjugate gradient minimization with restraints on heavy atoms of the protein and RNA, (2) heating from 0 to 310 K followed by 10 ps of MD with above restraints, (3) 1000 steps of minimization without restraints, (4) heating from 0 to 310 K followed by 100 ps of unrestrained equilibration. We used 1 fs MD integration steps for the equilibration phase and 2 fs for the 200 ns production phase. The final snapshots were visualized using PyMol[13].


1 Huang, F. Efficient incorporation of CoA, NAD and FAD into RNA by in vitro transcription. *Nucleic Acids Research* **31**, e8-e8, doi:10.1093/nar/gng008 (2003).

2 Rasband, W. S. ImageJ. *U. S. National Institutes of Health, Bethesda, Maryland, USA,* (1997-2018).

3 Datsenko, K. A. & Wanner, B. L. One-step inactivation of chromosomal genes in <em>Escherichia coli</em> K-12 using PCR products. *Proceedings of the National Academy of Sciences* **97**, 6640-6645, doi:10.1073/pnas.120163297 (2000).

4 Goh, G. B., Hodas, N. O. & Vishnu, A. Deep learning for computational chemistry. *Journal of Computational Chemistry* **38**, 1291-1307, doi:doi:10.1002/jcc.24764 (2017).

5 Alarcos, N., Cohen, B., Ziółek, M. & Douhal, A. Photochemistry and Photophysics in Silica-Based Materials: Ultrafast and Single Molecule Spectroscopy Observation. *Chemical Reviews* **117**, 13639-13720, doi:10.1021/acs.chemrev.7b00422 (2017).

6 Palermo, G., Miao, Y., Walker, R. C., Jinek, M. & McCammon, J. A. Striking Plasticity of CRISPR-Cas9 and Key Role of Non-target DNA, as Revealed by Molecular Simulations. *ACS Central Science* **2**, 756-763, doi:10.1021/acscentsci.6b00218 (2016).

7 Berendsen, H. J. C., Grigera, J. R. & Straatsma, T. P. The missing term in effective pair potentials. *The Journal of Physical Chemistry* **91**, 6269-6271, doi:10.1021/j100308a038 (1987).

8 Wang, J. *et al.* Twin Defect Derived Growth of Atomically Thin MoS2 Dendrites. *ACS Nano* **12**, 635-643, doi:10.1021/acsnano.7b07693 (2018).

9 Yang, Q. *et al.* Donor Engineering for NIR-II Molecular Fluorophores with Enhanced Fluorescent Performance. *Journal of the American Chemical Society* **140**, 1715-1724, doi:10.1021/jacs.7b10334 (2018).

10 Shupanov, R., Chertovich, A. & Kos, P. Micellar polymerization: Computer simulations by dissipative particle dynamics. *Journal of Computational Chemistry* **39**, 1275-1284, doi:doi:10.1002/jcc.25194 (2018).

11 Gaussian 09, Revision A.02 (2016).

12 .A. Case, R. M. B., D.S. Cerutti, T.E. Cheatham, III, T.A. Darden, R.E. Duke, T.J. Giese, H. Gohlke, A.W. Goetz, N. Homeyer, S. Izadi, P. Janowski, J. Kaus, A. Kovalenko, T.S. Lee, S. LeGrand, P. Li, C. Lin, T. Luchko, R. Luo, B. Madej, D. Mermelstein, K.M. Merz, G. Monard, H. Nguyen, H.T. Nguyen, I. Omelyan, A. Onufriev, D.R. Roe, A. Roitberg, C. Sagui, C.L. Simmerling, W.M. Botello-Smith, J. Swails, R.C. Walker, J. Wang, R.M. Wolf, X. Wu, L. Xiao and P.A. Kollman. AMBER. *University of California, San Francisco* (2016).

13 DeLano, W. L. Pymol: An open-source molecular graphics tool. *CCP4 Newsletter On Protein Crystallography* **40**, 82-92 (2002).
