## Extended data for "Dinucleoside polyphosphates act as 5’-RNA caps in *Escherichia coli*"

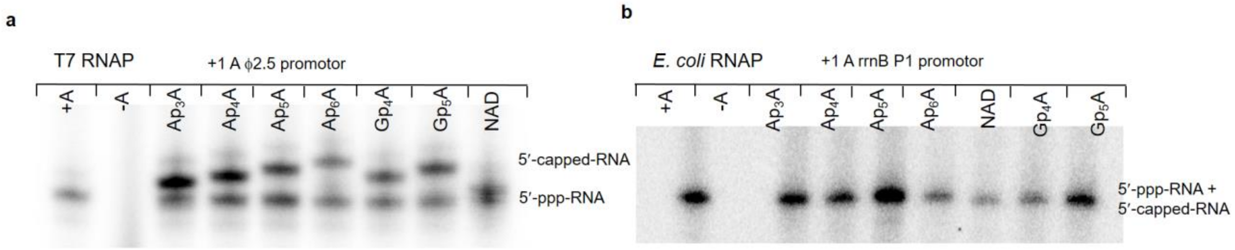


**c**

**+1 A ϕ2.5 promotor**

| conc. of cap [mM] | $Ap_3A$-RNA | $Ap_4A$-RNA | $Ap_5A$-RNA | $Ap_6A$-RNA | NAD |
|---|---|---|---|---|---|
| **0.2** | 16 ± 3.6 | 16 ± 2.9 | 18 ± 1.7 | 7 ± 1.7 | 13 ± 1.4 |
| **0.4** | 25 ± 2.6 | 23 ± 0.9 | 22 ± 2.5 | 15 ± 0.8 | 21 ± 2.5 |
| **1.0** | 37 ± 2.9 | 38 ± 3.1 | 38 ± 4.6 | 27 ± 1.7 | 33 ± 2.4 |
| **1.6** | 51 ± 6.0 | 48 ± 1.9 | 43 ± 2.9 | 36 ± 1.2 | 42 ± 2.6 |

**d**

**+1 G ϕ6.5 promotor**

| conc. of cap [mM] | $Gp_4G$-RNA | $Ap_4G$-RNA | $Ap_5G$-RNA |
|---|---|---|---|
| **0.2** | 22 ± 2.1 | 22 ± 0.9 | 21 ± 1.6 |
| **0.4** | 28 ± 2.1 | 30 ± 2.1 | 29 ± 3.7 |
| **1.0** | 44 ± 2.4 | 46 ± 2.9 | 42 ± 3.3 |
| **1.6** | 53 ± 2.1 | 55 ± 2.2 | 46 ± 5.2 |

*Extended Data Figure 1: In vitro* transcription with T7 and *E. coli* RNAP. a) *In vitro* transcription with T7 RNAP, template containing promotor Aϕ2.5 and various $Np_nNs$ or NAD as NCIN. b) *In vitro* transcription with *E. coli* RNAP, template containing promotor A rnnB P1 and various $Np_nNs$ or NAD as NCIN. c) Table of the amount of capped RNA (in %) compared to the uncapped 5'-ppp-RNA prepared by *in vitro* transcription with T7 RNAP in the presence of constant amount of ATP (1 mM) and increasing amount (0.2-1.6 mM) of $Np_nNs$ or NAD (caps) in the solution. The percentage was determined after PAGE analysis of the *in vitro* transcription using 35A or 35G template. The RNA was labelled by $\alpha$-$^{32}P$ GTP. The average values and their standard deviations were calculated from triplicates.

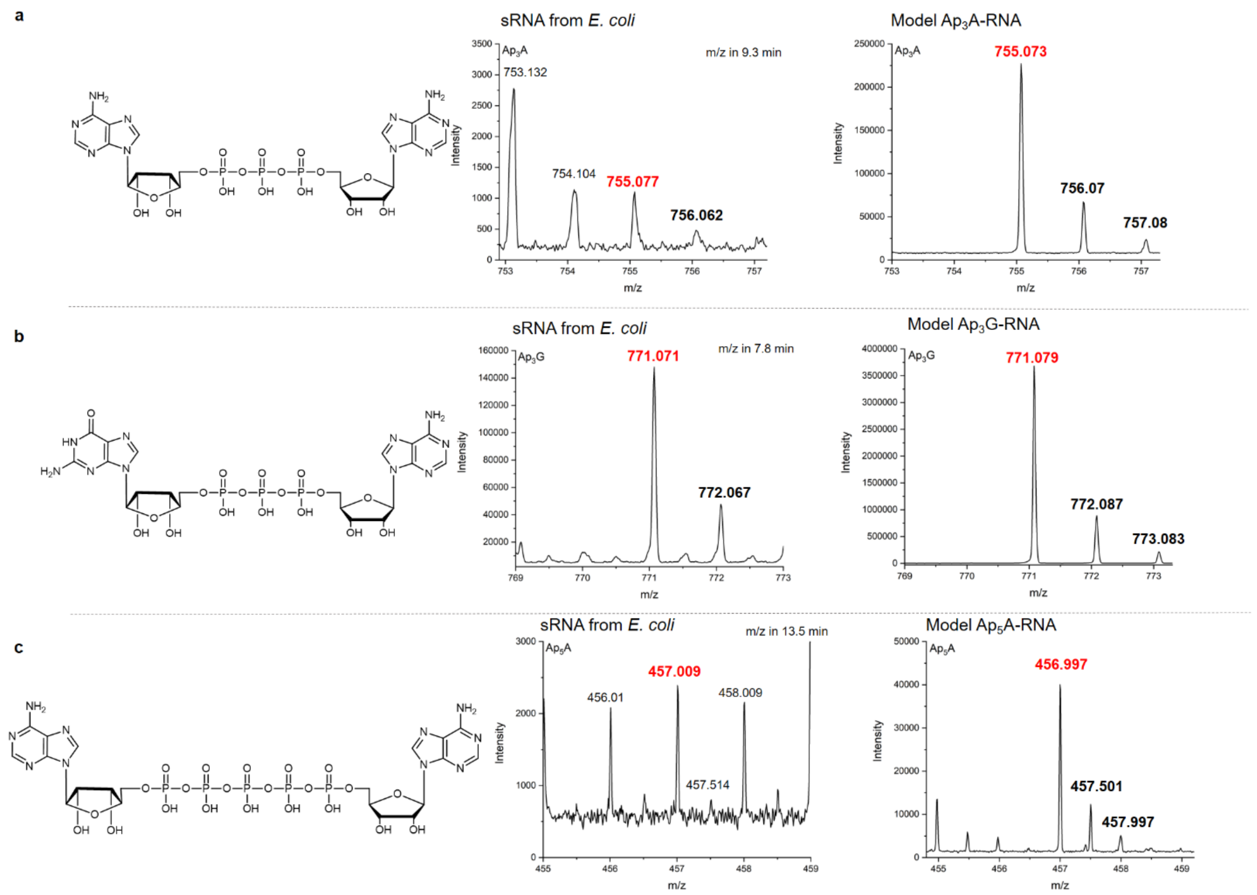


*Extended Data Figure 2:* LC-MS detection of naturally occurring Np$_n$N-RNA in *E. coli.* From the top structures of Ap$_3$A (**a**), Ap$_3$G (**b**) and Ap$_5$A (**c**) and comparison of observed MS spectra in RNA from *E. coli* and synthetic standards.

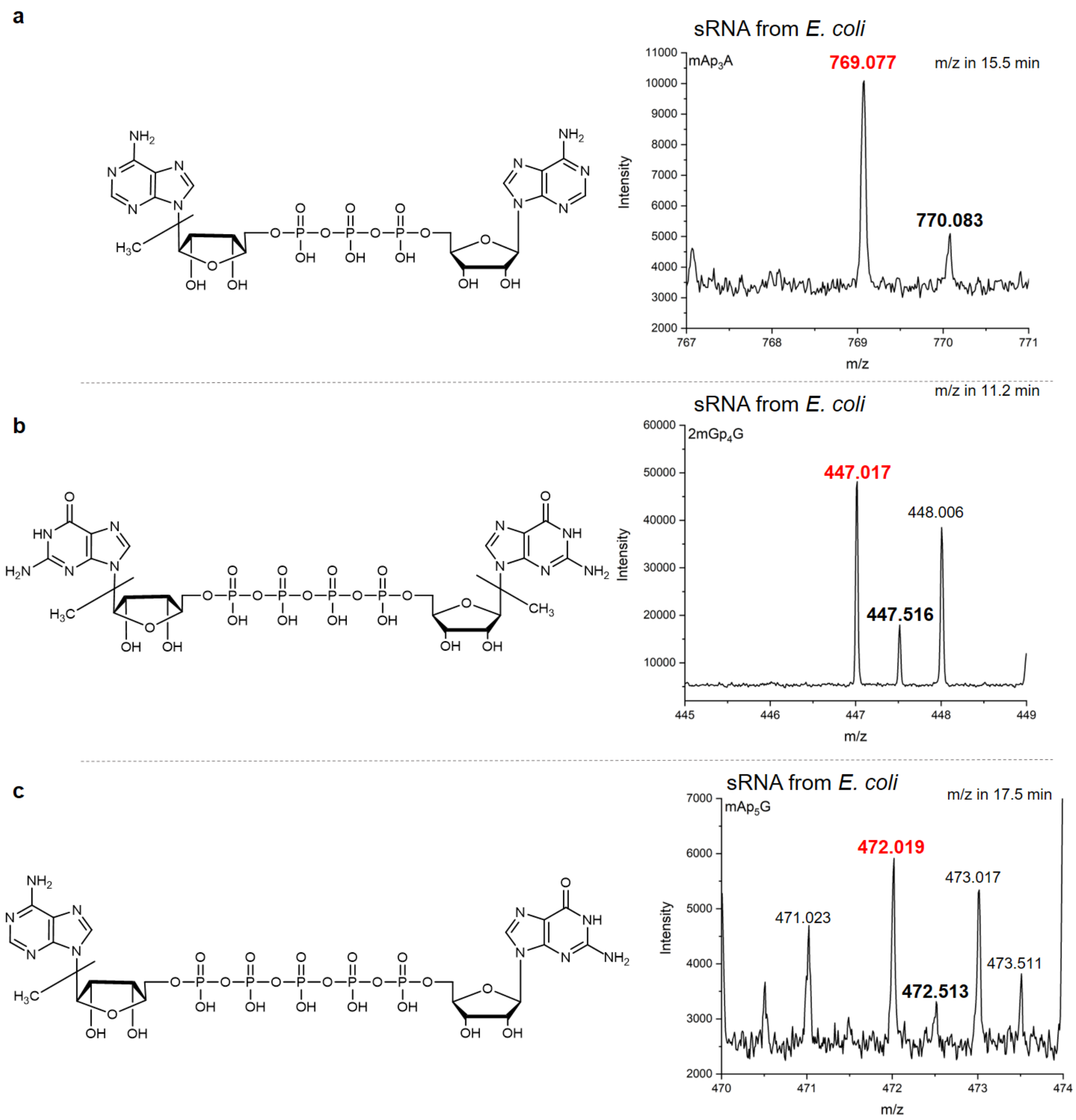


*Extended Data Figure 3:* LC-MS detection of naturally occurring Np*n*N-RNA in *E. coli.* From the top structures of $mAp_3A$ (**a**), $2mGp_4G$ (**b**) and $mAp_5G$ (**c**) and MS spectra of observed caps in RNA from *E. coli*.

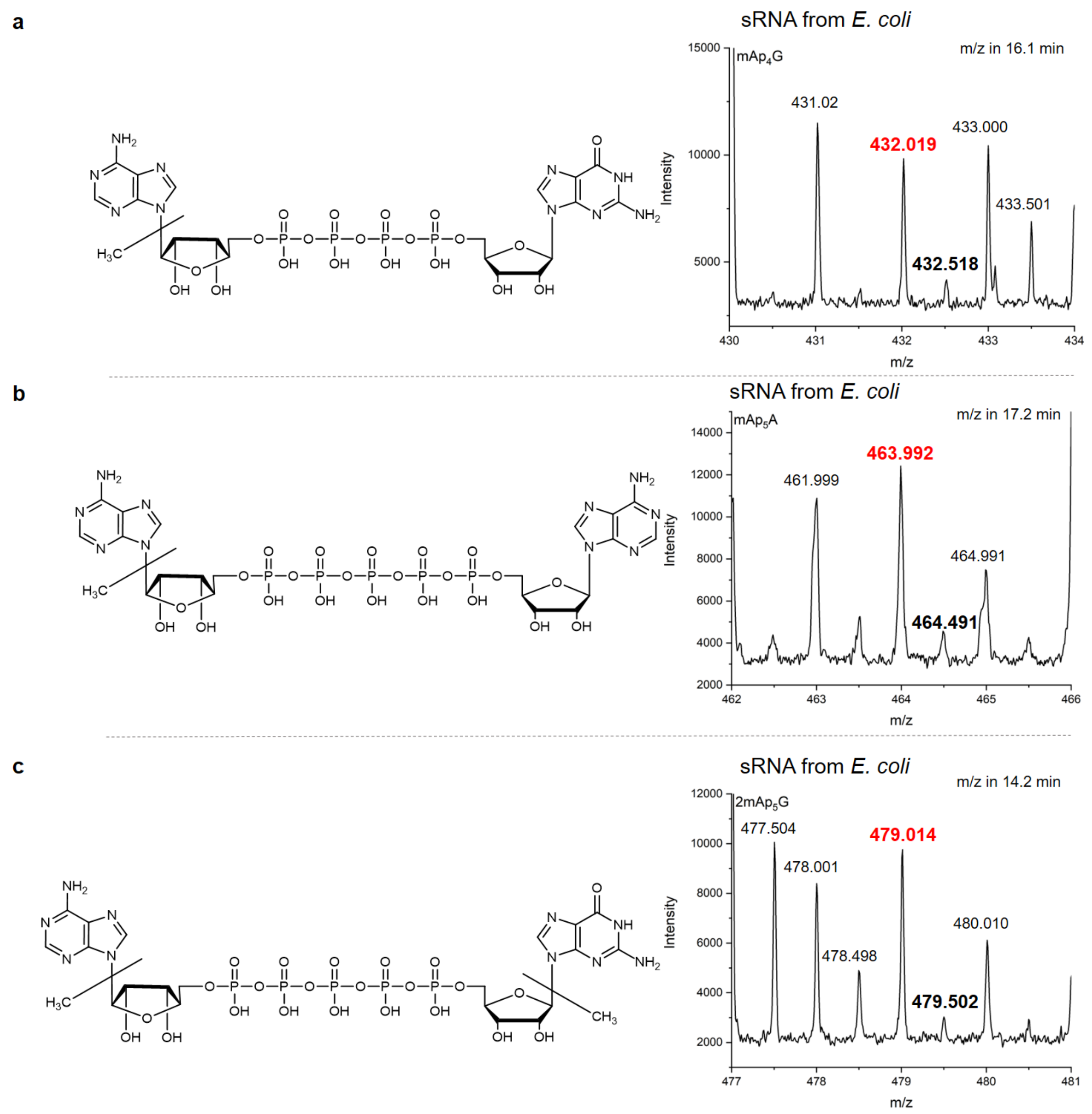


*Extended Data Figure 4:* LC-MS detection of naturally occurring Np$_n$N-RNA in *E. coli.* From the top structures of mAp$_4$G (**a**), mAp$_5$A (**b**) and 2mAp$_5$G (**c**) and MS spectra of observed caps in RNA from *E. coli*.

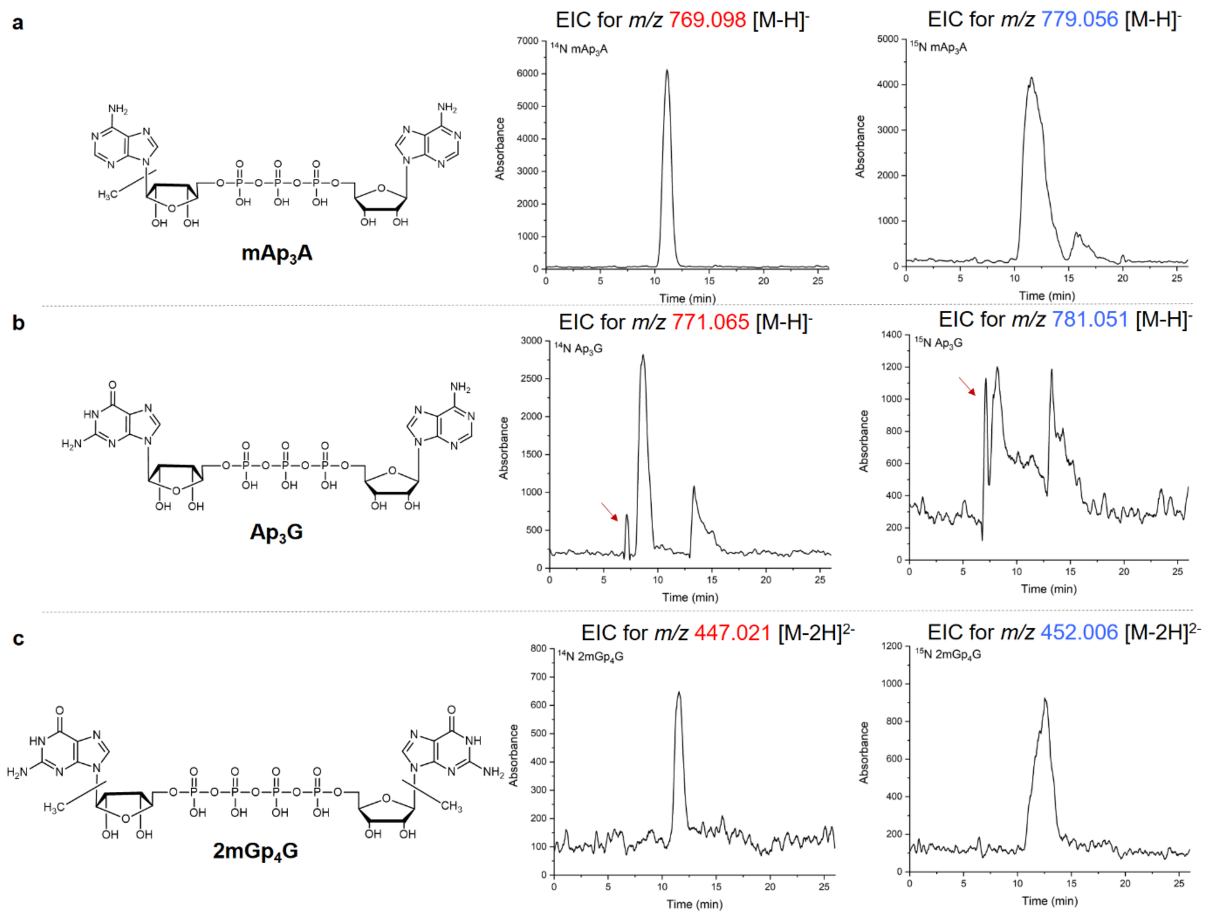


*Extended Data Figure 5:* Structures of different RNA caps and EIC for various *m/z* in RNA from *E. coli* growth in minimal media containing $^{14}N$ or $^{15}N$ of methyl-$Ap_3A$ (**a,** in $^{14}N$ *m/z* 769.085 was shifted to *m/z* 779.056 in $^{15}N$ in 11 min retention time), $Ap_3G$ (**b,** in $^{14}N$ *m/z* 771.069 was shifted to *m/z* 781.035 in $^{15}N$ in 7 min retention time), dimethyl-$Gp_4G$ (**c,** in $^{14}N$ *m/z* 447.021 was shifted to *m/z* 452.006 in $^{15}N$ in 12 min retention time).

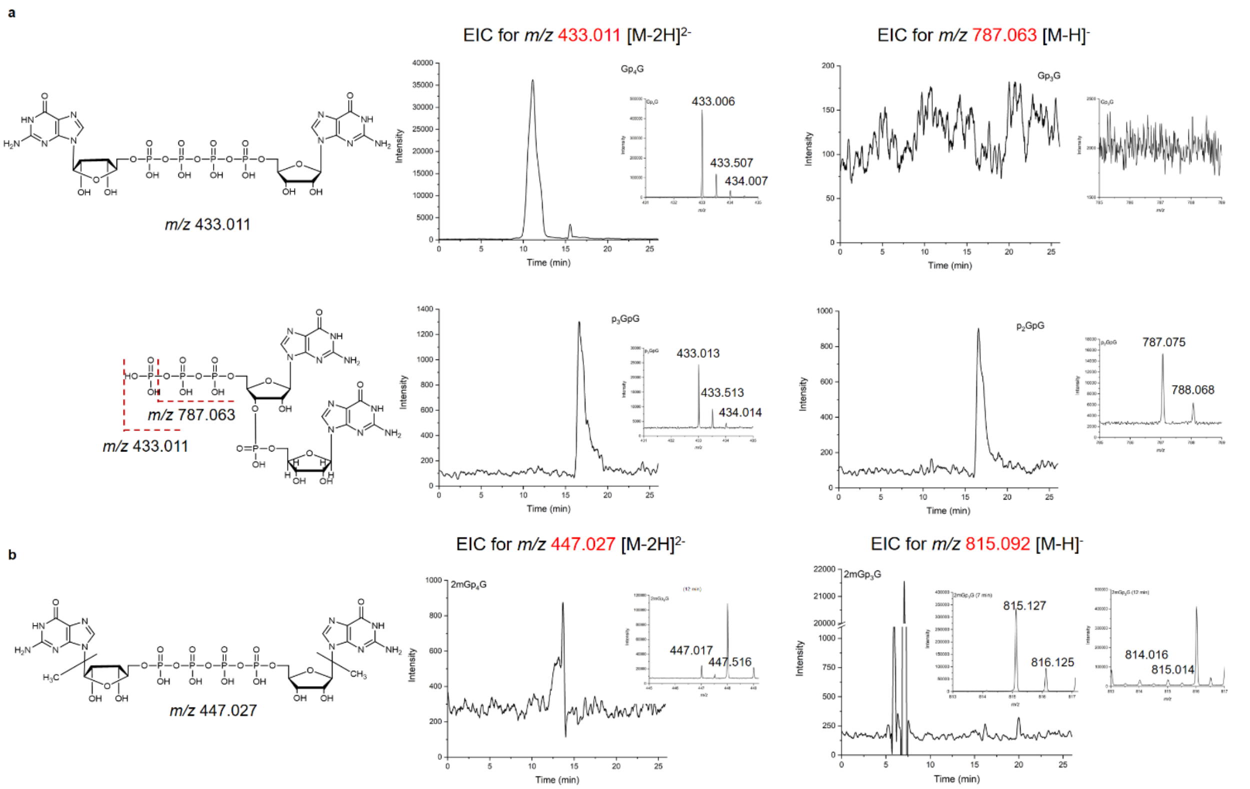


*Extended Data Figure 6:* Confirmation of 5'-5''polyphosphate chain. a) Fragmentation of synthetic standards. Structure and LC-MS analysis of model $Gp_4G$ and $p_3GpG$. Extracted Ion Chromatograms (EIC) for *m/z* 433.011 ($Gp_4G$ and $p_3GpG$, $[M-2H]^{2-}$) (middle) and for *m/z* 787.063 ($Gp_3G$ and $p_2GpG$, $[M-H]^-$) (right). Molecule $p_3GpG$ with external polyphosphate chain undergoes the cleavage under the ionization conditions and *m/z* 787.063 is observed. While $Gp_4G$ with internal polyphosphate chain stays under the same conditions intact. b) Structure and LC-MS analysis of detected $2mGp_4G$. EIC for *m/z* 447.027 ($2mGp_4G$. $[M-2H]^{2-}$) (middle) and for *m/z* 815.092 ($2mGp_3G$, $[M-H]^-$) (right). EIC is comparable with EIC of model $Gp_4G$ confirming the internal polyphosphate chain.

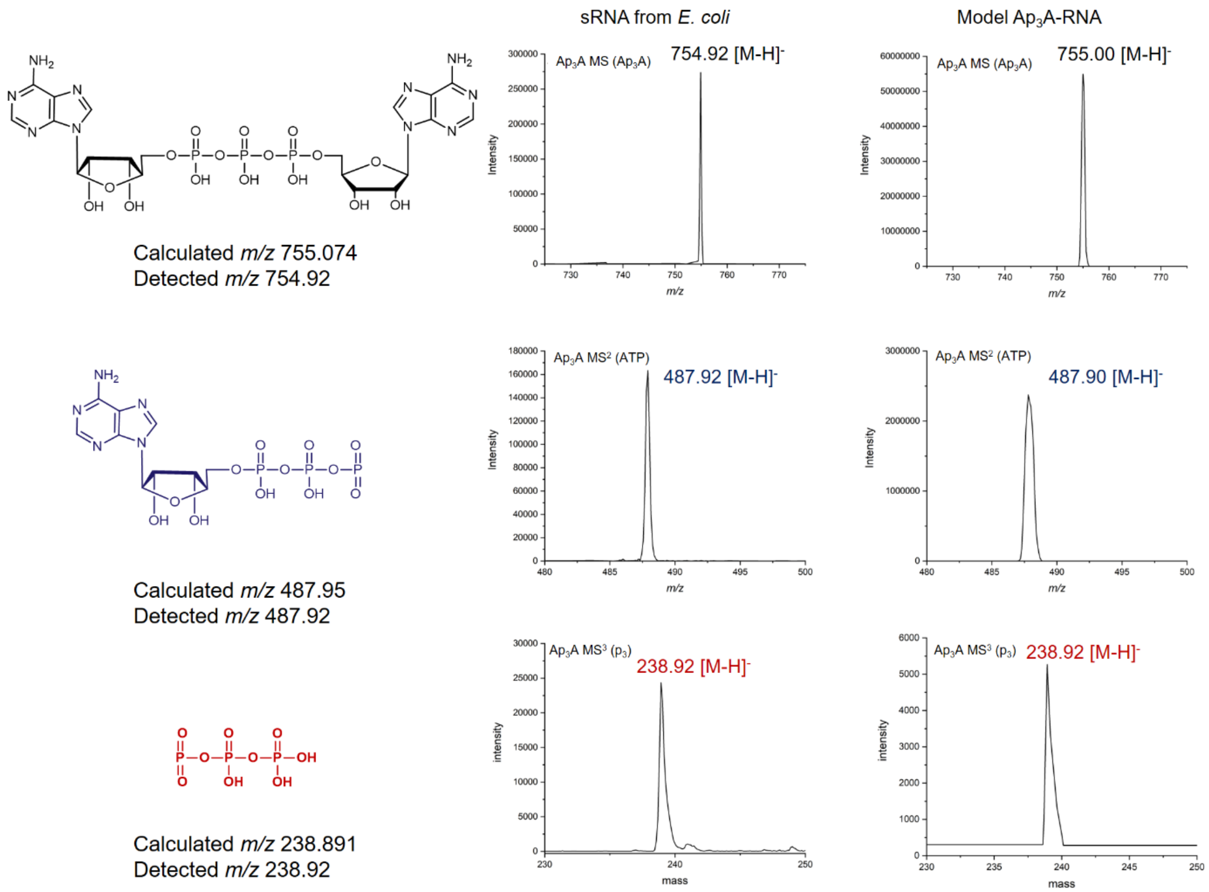


*Extended Data Figure 7:* The fragmentation behaviour of detected $Ap_3A$. $Ap_3A$ ion was fragmented and peak with *m/z* 487.8 (ATP, $[M\text{-}H]^-$) was selected for further fragmentation ($MS^3$). The *m/z* 238.92 $[M\text{-}H]^-$ corresponds to the intact polyphosphate chain. The same behaviour was observed for $Ap_3A$ standard (right) as for $Ap_3A$ coming from the analysed sample from RNA of *E. coli* (middle).

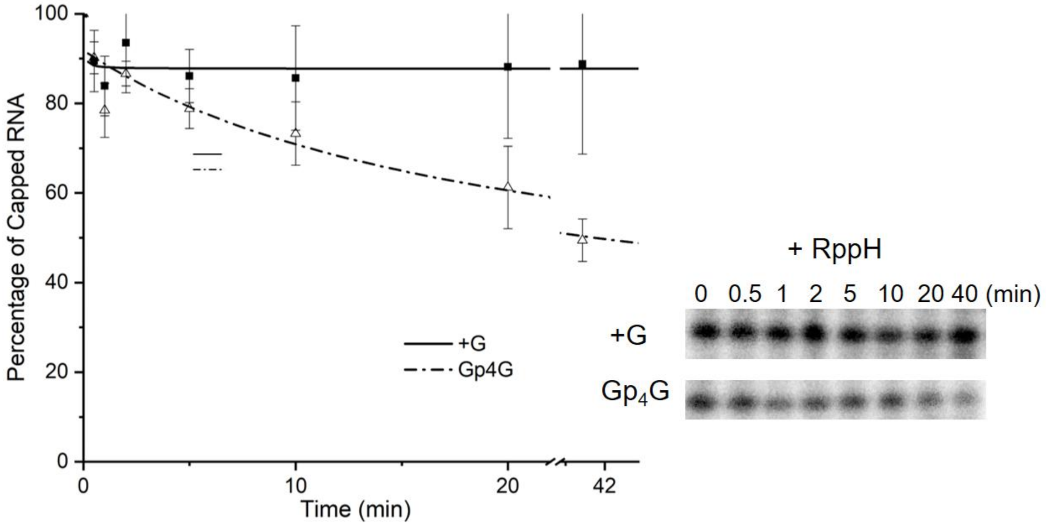


*Extended Data Figure 8:* Kinetic studies of RppH cleavage of $Gp_4G$-capped in comparison with pppG-RNA stopped after 30 s, 1, 2, 5, 10, 20 and 40 min and analysed by PAGE.

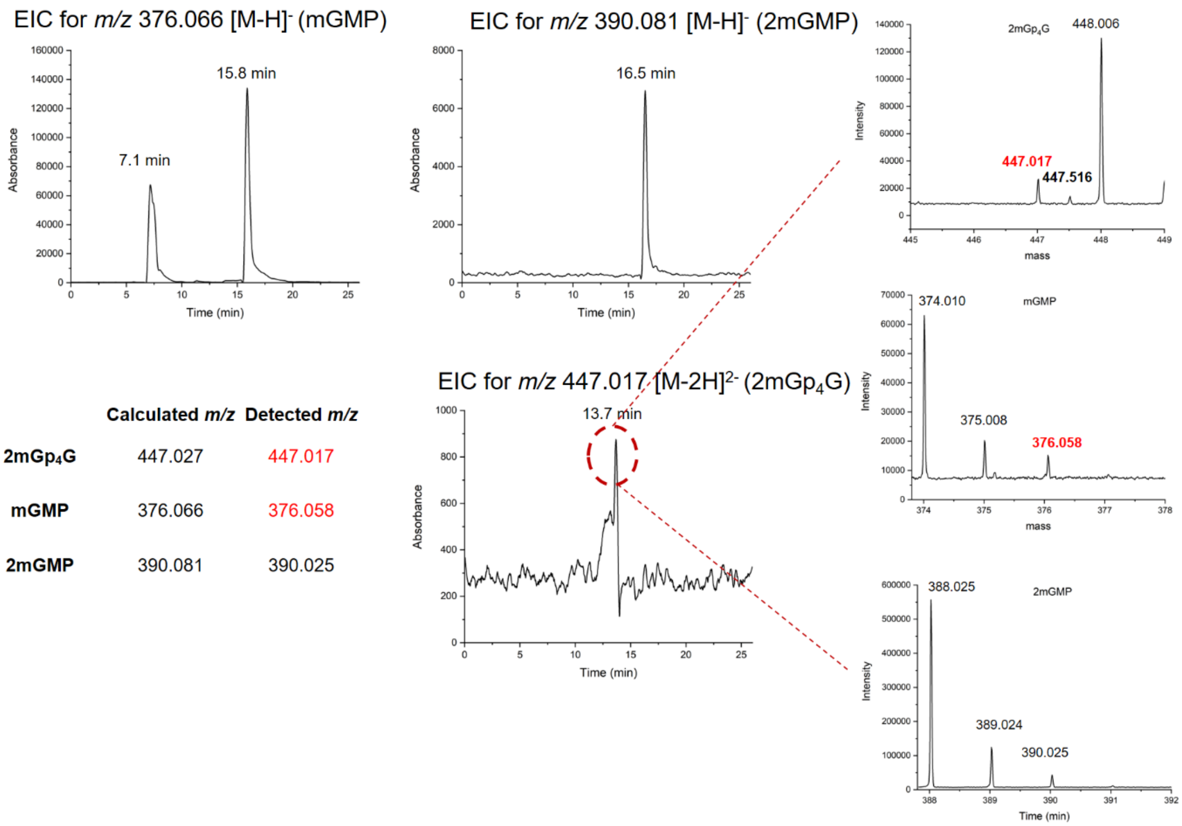


| | Calculated *m/z* | Detected *m/z* |
|---|---|---|
| **2mGp₄G** | 447.027 | 447.017 |
| **mGMP** | 376.066 | 376.058 |
| **2mGMP** | 390.081 | 390.025 |

*Extended Data Figure 9:* Distribution of methyl groups in detected 2mGp$_4$G. EIC for *m/z* 376.066 (mGMP, [M-H]$^-$) and *m/z* 390.081 (2mGMP, [M-H]$^-$) in sRNA from *E. coli*. EIC for *m/z* 447.017 (2mGp$_4$G, [M-2H]$^{2-}$) and MS spectra for 13.7 min. The MS spectra confirmed the presence of monomethylated GMP (*m/z* 376.058, [M-H]$^-$) as fragment coming from 2mGp$_4$G. The calculated *m/z* 390.081 corresponding to dimethylated GMP was not detected in time 13.7 min. Detected ion of m/z 390.025 is not 2mGMP since it is isotopic *m/z* of *m/z* 388.025 and the error of the measurement is more than 140 ppm. The average error of the measurement is 25 ppm.
